# Genome shuffling in a globalized bacterial plant pathogen: Recombination-mediated evolution in *Xanthomonas euvesicatoria* and *X. perforans*

**DOI:** 10.1101/180901

**Authors:** Mustafa O. Jibrin, Neha Potnis, Sujan Timilsina, Gerald V. Minsavage, Gary E. Vallad, Pamela D. Roberts, Jeffrey B. Jones, Erica M. Goss

## Abstract

Bacterial recombination and clonality underly the evolution and epidemiology of pathogenic lineages as well as their cosmopolitan spread. While the spread of stable clonal bacterial pathotypes drives disease epidemics, recombination leads to the evolution of new bacterial lineages. Recombinant lineages of plant bacterial pathogens are typically associated with colonization of novel hosts and emergence of new diseases. Here, we show that recombination between evolutionarily and phenotypically distinct plant pathogenic lineages has generated new recombinant lineages with unique combinations of pathogenicity and virulence factors. *X. euvesicatoria* (*Xe*) and *X. perforans* (*Xp*) are two closely related monophyletic species causing bacterial spot disease on tomato and pepper worldwide. We sequenced the genomes of strains representing populations on tomato in Nigeria and found shuffling of secretion systems and effectors such that these strains contain genes from both *Xe* and *Xp*. Multiple strains, from populations in Nigeria, Italy, and Florida, USA, exhibited extensive genomewide homologous recombination and both species exhibited dynamic open pangenomes. Our results show that recombination is generating new lineages of bacterial spot pathogens on tomato with consequences for disease management strategies.

**Importance:** The *Xanthomonas* pathogens that cause bacterial spot of tomato and pepper have been model systems for plant-microbe interactions. Two of these pathogens, *X. euvesicatoria* and *X. perforans*, are very closely related. Genome sequences of bacterial spot field strains from Nigeria, Italy, and the United States showed varying levels of homologous recombination that changed the amino sequence of effectors, secretion systems, and other proteins. This shuffling of genome content occurred between *X. euvesicatoria* and *X. perforans*, while a Nigerian lineage also contained the lipopolysaccharide cluster of a distantly related *Xanthomonas* species. Gene content varied among strains and the affected genes are important in the establishment of disease, therefore our findings point to global variation in the host-pathogen interaction driven by gene exchange among evolutionarily distinct lineages.

## Introduction

While what defines a bacterial species remains contentious, our understanding the evolution of bacterial lineages is dependent on the elucidation of the various forms of adaptive divergence they experience (1-3). Bacterial population structure may change over time, such that an historically clonal bacterial population can experience recombination to the point of panmixis and return to clonality as ecologically stable forms emerge and increase in frequency by positive selection (4-6). Processes that lead to ecologically stable populations include the cessation of panmixis accompanied by continued accumulation of mutations in what is now described as the ecotype model (7-8). Clonality maintains adaptive gene complexes and produces the epidemic population structure described by Maynard Smith and colleagues (4). This conceptual model has been used to describe the emergence, spread and cosmopolitan fitness of virulent clones of the O1 and O139 serotypes of *Vibrio cholera*, and evolutionary processes leading to the divergence of clades of *Bacillus simplex* and *Mycobacterium tuberculosis* (6; 9-11). Among plant pathogens, populations of rapidly spreading bacteria such as *Pseudomonas syringae* pv. *actinidae* and *P. syringae* pv. *aesculi* causing the kiwifruit canker epidemic in Europe/New Zealand and bleeding canker disease of horse chestnut in Europe, respectively, are viewed as emergent clonal pathotypes that have colonized a new ecological niche (12-13).

In the ecotype model, non-homologous and homologous recombination are major drivers of evolution in bacterial populations (14-17; 6). Horizontal gene transfer (HGT, also called non-homologous recombination or lateral gene transfer), occurs between and within species (18). Transformation, transduction and plasmid-mediated gene transfer are all mechanisms through which HGT is known to occur in bacteria (19). The rate of HGT can be high and produces substantial variation in gene content even among members of a taxonomic group (20, 14; 21). Recombination can spread beneficial mutations (otherwise at risk of elimination due to clonal interference) and bring together independently evolved beneficial mutations (15). The result is that DNA sequences of new bacterial lineages are often mosaics of several closely related bacterial sequences (22). While HGT involves the direct transfer and receipt of genes between strains, homologous recombination requires some level of sequence homology. Previously, the prevailing model was that a high degree of sequence identity was important and that the efficiency of recombination decreased greatly as identity decreased (23). However, it has been shown that the level of sequence identity does not need to be constant across the gene length, rather, it is most important within minimum efficiently processed segments (MEPS) which are present in the inserted DNA’s flanking regions (16). In fact, the regions between MEPS may not share homology. The need for some level of homology means that recombination is highest among close relatives and across regions of the ‘core’ genome where homology is retained (24, 25).

Advancements in genome sequencing technology has led to genome-based analyses of bacterial recombination. These studies have shown the importance of recombination in the evolution of major bacterial pathogens of humans (26, 24-25, 27-28). Genomic studies on the relative role of recombination in plant pathogenic bacterial evolution remain limited. Recombination has played a role in host shifts by xanthomonads infecting Brassicaceae and citrus (29-30). Adaptation to agricultural crops by *Pseudomonas* pathogens has also been shown to be influenced by recombination (31). Plant disease management strategies could be aided by improved understanding of the contribution of recombination to plant bacterial pathogen evolution.

The genus *Xanthomonas*, which consists of more than twenty-seven described pathogenic species infecting over four hundred plant hosts, is one of the most important groups of bacteria causing plant disease (32). Bacterial spot of tomato and pepper is found around the world and is caused by four *Xanthomonas* species: *X. euvesicatoria* (*Xe*), *X. vesicatoria* (*Xv*), *X. perforans* (*Xp*) and *X. gardneri* (*Xg*) (33-35). Strains belonging to *Xe* and *Xp* are closely related (Potnis *et al.,* 2011) and may comprise a single species (36). Yet, *Xe* and *Xp* possess different sets of secreted effectors that elicit hypersensitive reactions on tomato differential lines (37, 35). Type III secretion effectors (T3SEs) are essential virulence factors that are secreted through the highly conserved type III secretion system (T3SS) and translocated into host plants where they interfere with host immunity (38). Within *Xp*, MLSA and core genome analyses have shown distinct evolutionary groups: 1A, 1B and 2 (39-40). A core genome phylogeny of *Xp* strains from Florida showed that Group 1A formed a monophyletic lineage of strains isolated in 2012. The reference strain for *Xp*, Xp91-118, represented one of multiple lineages in group 1B together with strains from 2006. Group 2 strains formed another distinct lineage and represented the bulk of all sequenced strains, and included strains from 2006, 2010, 2012 and 2013 (40). Irrespective of these groupings, there are effectors that have been consistently associated with either *Xe* or *Xp* and are described as *Xe-*specific and *Xp-*specific effectors (35, 40). Other studies have found unique bacterial spot strains in Europe and Africa, suggesting the possibility of additional evolutionarily distinct lineages (41-44).

We previously reported the occurrence in Nigeria of atypical bacterial spot of tomato strains (represented by strain NI1). These strains, collected in 2014, were identified as *Xp* because differential reactions on tomato genotypes indicated the presence of the *Xp* effector protein AvrXv3, and the highly conserved gene *hrcN* showed 100% sequence identity to all previously isolated *Xp* strains (44). However, multi-locus sequence analysis using six housekeeping genes clustered these Nigerian strains with *Xe*. Specifically, the *gltA*, *lacF*, and *gapA* sequences were identical or nearly identical to those of *Xe*, yet genes *fusA* and *gyrB* were identical to alleles found in *Xp* group 1 strains and the *lepA* allele was distinct from both *Xe* and *Xp* (39). Strains like NI1 were isolated from the same tomato field in 2015, but we also identified a second group of strains. Differential reactions indicated that these strains contained the *Xe* effector protein AvrRxv. However, the *hrcN* gene sequence placed the strains in *Xp*. This was surprising because no *Xp* field strain has been reported to contain *avrRxv*. The presence of this gene was confirmed in representative strain NI38 by Sanger sequencing (Jibrin, Roberts, Jones and Minsavage, *unpublished*). In summary, genotypic and phenotypic tests typically used to assign bacterial spot strains to species could not assign the Nigerian strains to a *Xanthomonas* species due to conflicting race, MLSA, and/or *hrcN* data.

The objective of this study was to examine the evolutionary processes that led to the conflicting assignments of the Nigerian strains. We hypothesized that recombination had generated lineages of bacterial spot strains with unique T3SS and T3SE gene profiles. We are interested in T3SS effectors, because they are often targets of resistance breeding and are used to monitor pathogen population shifts (37). Representatives of the Nigerian strains, NI1 and NI38, were sequenced and compared with 63 previously sequenced strains, focusing mostly but not exclusively on *Xe* and *Xp.* Our results show that recombination is driving the evolution of new lineages of bacterial spot pathogens and thereby affecting management decisions.

## Results

### Average nucletide identity distinguishes NI1 from *Xe* and *Xp*

Whole genome sequences were obtained for NI1 and NI38 as representative strains of Nigerian bacterial spot populations. Average nucleotide identity (ANI) statistics showed >99% identity among strains within species (within *Xp* and within *Xe*) and <99% identity in *Xp* to *Xe* comparisons (Table 1; Supplementary File 1). In contrast, the NI1 strain showed only 98.9% and 98.5% identity in pairwise comparisons to *Xp* and *Xe* reference genomes, respectively (Table 1) and <99% identity to all other *Xp* and *Xe* strains (Supplementary File 1). The second strain representing Nigerian populations, NI38, showed ANIs typical of an *Xe* strain: ANIs were >99% in comparisons against *Xe* genomes and <99% against *Xp* genomes. The ANI results also showed that strains belonging to *Xp* group 1A had more conserved ANI (≥99.97) while more variability was found within groups 1B and 2 (Supplementary File 1).

**Table 1.** Average nucleotide identities (ANIb) of genomes of Nigerian, *X. perforans* and *X. euvesicatoria* strains as compared to reference genomes. For all pairwise comparisons, see Supplementary File 1.

| Species/Groups | ANi vs. Xp91-118 | ANi vs. Xe85-10 |
| --- | --- | --- |
| *Nigerian strains* |  |  |
| NI1 | 98.92 | 98.58 |
| NI38 | 98.48 | 99.79 |
| *X. perforans* |  |  |
| Xp91-118 | 100 |  |
| Group 1A strains | 99.87-99.90 | 98.47-98.50 |
| Group 1B strains | 99.70-99.86 | 98.36-98.55 |
| Group 2 strains | 99.67-99.78 | 98.44-98.62 |
| Xp17-12 |  |  |
| Xp4P1S2 | 99.59 | 98.44 |
| *X. euvesicatoria* |  |  |
| Xe85-10 | 98.48 | 100 |
| Other *Xe* strains | 98.32-98.44 | 99.66-99.95 |

### *Xe* and *Xp* have open pangenomes

Analysis of the *Xe* and *Xp* pan and core genomes showed heterogeneity in genome composition. We observed a gradual decrease in core genome size with the addition of strains, and a bimodal distribution in core genome size depending on the genomes re-sampled (Supplementary Figure S1). Likewise, the pangenome size was highly dependent on the strains sampled. For *Xe*, addition of the NI38 genome produced only a small increase in pangenome size (Supplementary Figure S1). For *Xp*, addition of the Italian strain Xp_4P1S2 and NI1 produced a noticeable uptick in the size of the *Xp* pangenome. The increasing size of the pangenomes as more strains were sampled indicates open pangenomes for both *Xe* and *Xp*.

To examine the differences in gene content between NI1, NI38 and other *Xp* and *Xe* strains, we produced a pangenome tree using Gower’s similarity coefficient, which is 0 when genomes have identical gene content and increases up to a maximum of 1 with increasing dissimilarity. The Gower’s coefficient values clearly showed NI1 to be intermediate between *Xe* and *Xp* (Figure 1), which is in agreement with the ANI results. NI38 clustered with other *Xe* strains.

### Phylogenetic reconstruction groups NI1 with *Xp* and NI38 with *Xe*

The maximum likelihood phylogeny for concatenated core genes showed *Xe* and *Xp* as diverged monophyletic groups, consistent with previous studies (Figure 2; 39-40). NI1 and 4P1S2 formed distinct lineages of *Xp*, while NI38 is an *Xe* lineage. NI1 is substantially diverged from the other *Xp* strains, but the genetic distance between NI1 and other *Xp* strains was not as great as the interspecific distance between *Xp* and *Xe*.

### Extensive recombination in lineages of *Xp*

ClonalFrameML analyses showed different rates of homologous recombination in *Xe* and *Xp* (Table 2). The ratios of recombination to mutation for *Xe* showed that homologous recombination of imported DNA occurred at less than half the rate of mutation (R/θ) across all *Xe* strains. For *Xp*, estimated rates of recombination to mutation were dependent on the population used for analysis. For all *Xp* strains, recombination and mutation approached the same rate (R/θ=0.85). When strains from the Florida population were considered exclusively (NI1 and 4P1S2 excluded), recombination reached 1.5 times the rate of mutation. In all cases, the overall impact of recombination on nucleotide variation was greater than that of mutation (R/m > 1).

**Table 2.** Rates of recombination as calculated using ClonalFrameML^1^.

| Xanthomonas<br>population | R/θ | δ | ν | R/m | Tr/Tv |
| --- | --- | --- | --- | --- | --- |
| All strains | 0.49 | 292 | 0.03 | 3.82 | 5.36 |
| All *Xp* (+NI1) | 0.85 | 394 | 0.02 | 8.47 | 4.92 |
| Florida *Xp*<sup>#</sup> | 1.54 | 696 | 0.02 | 26.23 | 3.47 |
| All *Xe* | 0.43 | 409 | 0.028 | 5.06 | 5.35 |
| *Xe* w/out NI38 | 0.36 | 731 | 0.015 | 4.02 | 4.02 |

<sup>1</sup>Values were rounded for presentation. R/θ is the rate of recombination to mutation; δ is the size of the recombined fragment in bp; ν is the probability of recombination per site; R/m is the relative effect of recombination to mutation (or ratio of per base pair substitution by recombination to mutation, R/m= R/θ x ν x δ); Tr/Tv is the transition/transversion ratio used as input in the analysis.
<sup>#</sup> NI1 (Nigeria) and 4P1S2 (Italy) were excluded.

The extensive homologous recombination in lineages of *Xe* and *Xp* is visualized in Figure 3. The number of recombination events is generally high in ancestral strains of each lineage, with a mostly clonal genealogy within each lineage. Strain NI1 had a particularly high number of inferred recombination events, and large chromosomal replacements, shown in Figure 3 by long continuous dark blue lines. Strains NI38, 4P1S2, Xp17-12, Xp5-6 and Xp4-20, which represent independent lineages, also had high numbers of recombination events. Regions of homoplasy were more common in strains of *Xe* than strains of *Xp,* suggesting that there may be more undetected recombination events in *Xe.*

### NI1 LPS cluster resembles the LPS cluster of X. translucens pv. translucens

Lipopolysaccharides are highly antigenic and often act as pathogen associated molecular patterns, virulence factors and defense response elicitors (45-47). The lipopolysaccharide (LPS) gene clusters in the genus *Xanthomonas* usually consist of 15 genes and are flanked by two conserved housekeeping genes, cystathionine gamma lyase (*metB)* and electron transport flavoprotein *(etfA)* (48). A previous comparison of the LPS clusters of bacterial spot xanthomonads using reference strains showed that *Xp* has a unique LPS cluster while *Xe, Xv* and *Xg* have similar clusters (35). We found high gene conservation and little variation among *Xp* strains (Figure 4b). In contrast, the NI1 LPS cluster contains several additional genes. For example, NI1 has two glycosyltransferase genes which belong to a cluster of orthologous genes (COG1216, GT2 family) that are absent in other *Xp* LPS clusters. BLAST searches of genes unique to the NI1 LPS cluster revealed similarity with the LPS cluster of *Xanthomonas translucens pv. translucens* (*Xtt)*, causal agent of bacterial wilt (also known as black chaff) of barley (*Hordeum vulgare*). Out of two available *Xtt* genome sequences on the IMG database, the NI1 LPS cluster was most similar to the *Xtt* type strain, DSM 18974, which was isolated from barley in the United States (49; Figure 4a).

### Recombination affected type 3 and type 4 secretion system genes

The Type 3 Secretion System (T3SS) is a delivery pathway for secreted effectors and avirulence proteins, which are effectors that are recognized by the plant thus triggering a resistance response. NI1 generally contains the expected *Xp* alleles for T3SS genes, but the *Xp* strain 4P1S2 contained a mixture of *Xe* and *Xp* alleles (Table 3). NI38 also exhibited a mix of *Xe* and *Xp* T3SS alleles (Table 3; Supplementary Table S3).

**Table 3.**
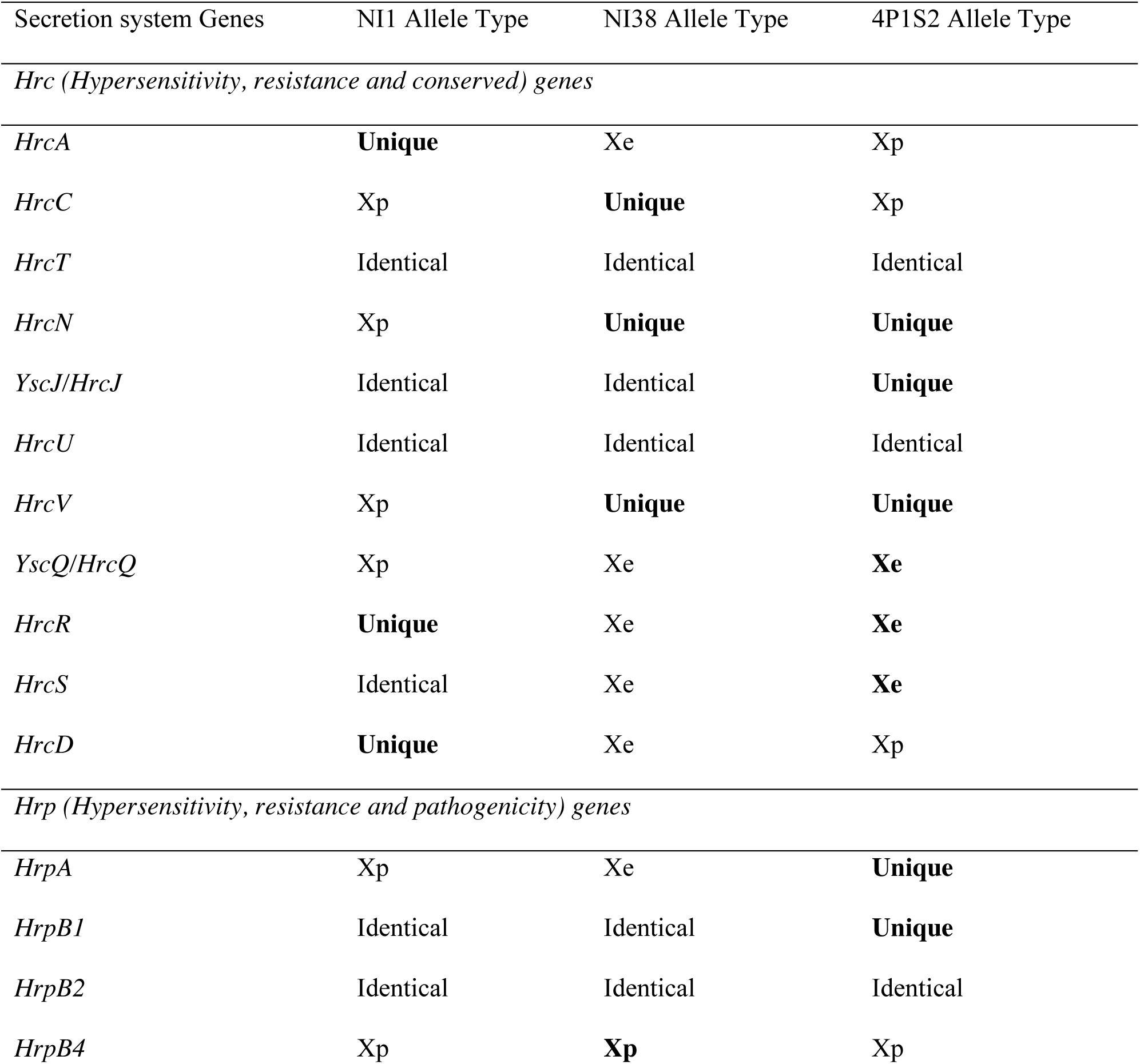
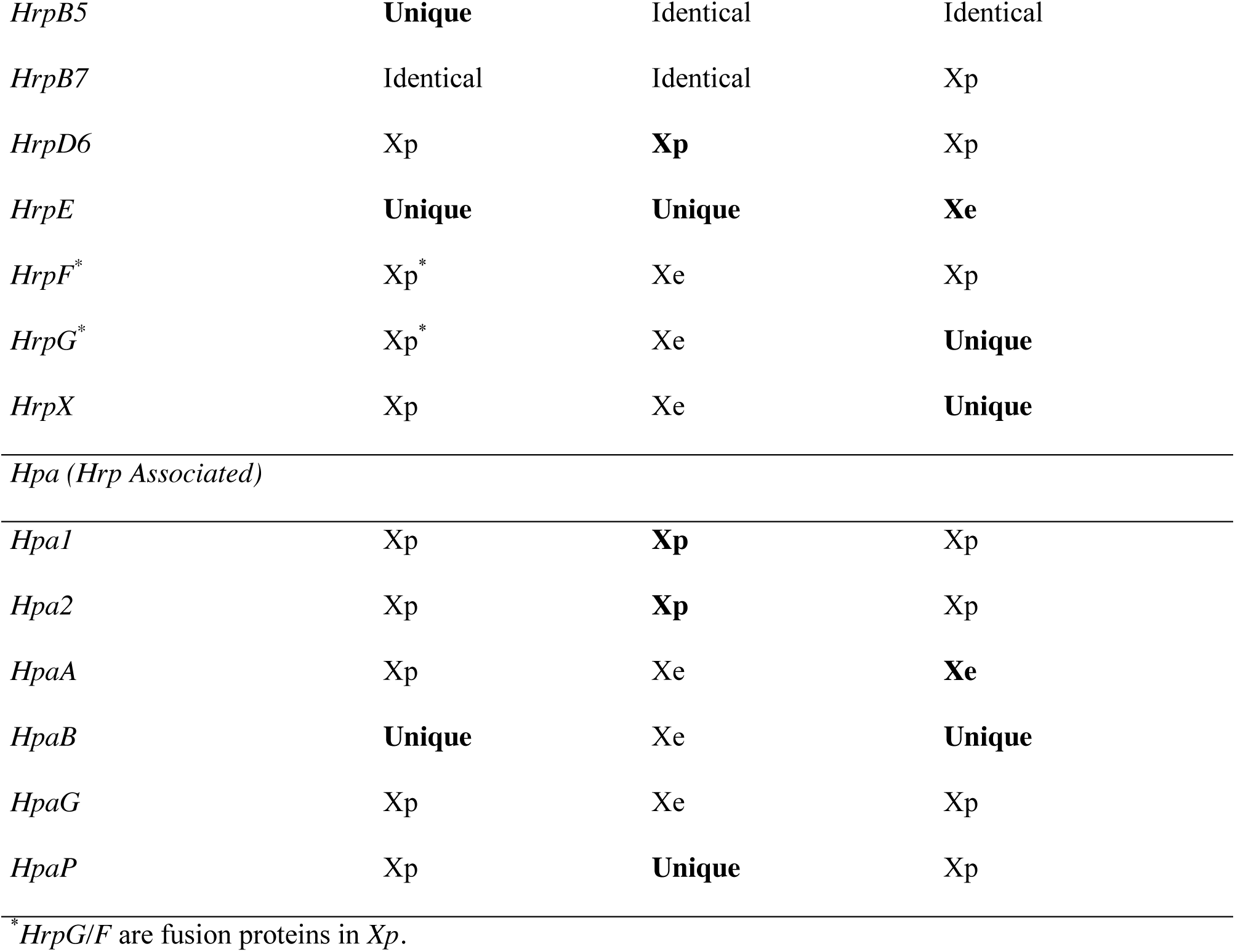
Allelic assignments of NI1, NI38 and 4P1S2 Type 3 Secretion System T3SS genes relative to *Xe* and *Xp* reference strains. Allele types are based on nucleotide sequence identity. ‘Identical’ indicates that *Xe* and *Xp* alleles were identical to each other, thus cannot be distinguished. ‘Unique’ indicates alleles not found in the reference strains. Allelic assignments based amino sequence identity are shown in Supplementary Table S3, because in some cases the nucleotide variation shown does not lead to variation in the protein sequence.

Two Type 4 Secretion Systems (T4SS) were reported in bacterial spot xanthomonads, the Vir and Dot/Icm systems (35). The reference strain for *Xe* has both systems and the type strain of *Xp* has only the Vir system. The T4SS is important for horizontal gene transfer between bacteria and delivery of effectors into hosts. Both types of T4SS were found in all *Xe* strains, including NI38, and also in NI1. With the exception of strain 4P1S2, all other *Xp* strains have a complete Vir system and only the *FimT* and *PilC* genes of the Dot/Icm system, lacking *PilE*, *PilV*, *PilW*, *PilX* and *PilY1* (Supplementary Table S4). Strain 4P1S2 is unique among these in having the *PilE* gene.

We did not identify variation in gene composition in the T2SS, T5SS or T6SS among strains. Both NI1 and NI38 had the type 1 and 3 T6SS clusters that were previously identified in bacterial spot xanthomonads (35).

### NI1 and NI38 contain a mix of *Xe*-specific and *Xp*-specific effectors

The mix of genes in the T3SS of NI1 and NI38 suggested the possibility of variability in T3SS effectors, which are often primary targets of resistance breeding (35, 40). The NI1 genome possesses all 11 previously described genes for core effectors, as well as *xopE2*, which was recently added to the core effectors of bacterial spot xanthomonads (Supplementary Table S5; 35, 40). Among the groups of effectors that are shared by *Xe* and *Xp* but absent in other bacterial spot xanthomonads, NI1 possesses all but *xopF2* (Table 4). Two of the four copies of *xopP* in NI1, together with *xopAK*, have 100% amino acid identity with the corresponding *Xp* effectors, but the *xopI* effector has 100% amino acid identity to the *Xe* allele of *xopI*. Some effectors have been previously described as species-specific among the bacterial spot xanthomonads. NI1 has *xopAJ* specific to *Xe*; *xopC2*, *xopAE* and *xopAF* specific to *Xp*; and *xopAQ* and *xvrHah1* specific to *X. gardneri* (*Xg*). While the *xopAJ* of NI1 shares 98.7% similarity to the *Xe* allele, it is 100% identical to a copy of *xopAJ* found in *X. axonopodis pv. poinsenttiicola*. NI1 lacked *xopJ4* which was found to be conserved among *Xp* strains in Florida (39). Consistent with this finding, NI1 did not cause a hypersensitive response on tomato plants containing the *RXopJ4* resistance gene (Figure 5). Strain 4P1S2 is similar to NI1 in that it contains *avrHah1* and *xopAQ*, however, it is similar to other *Xp* in having *xopJ4*.

**Table 4.**
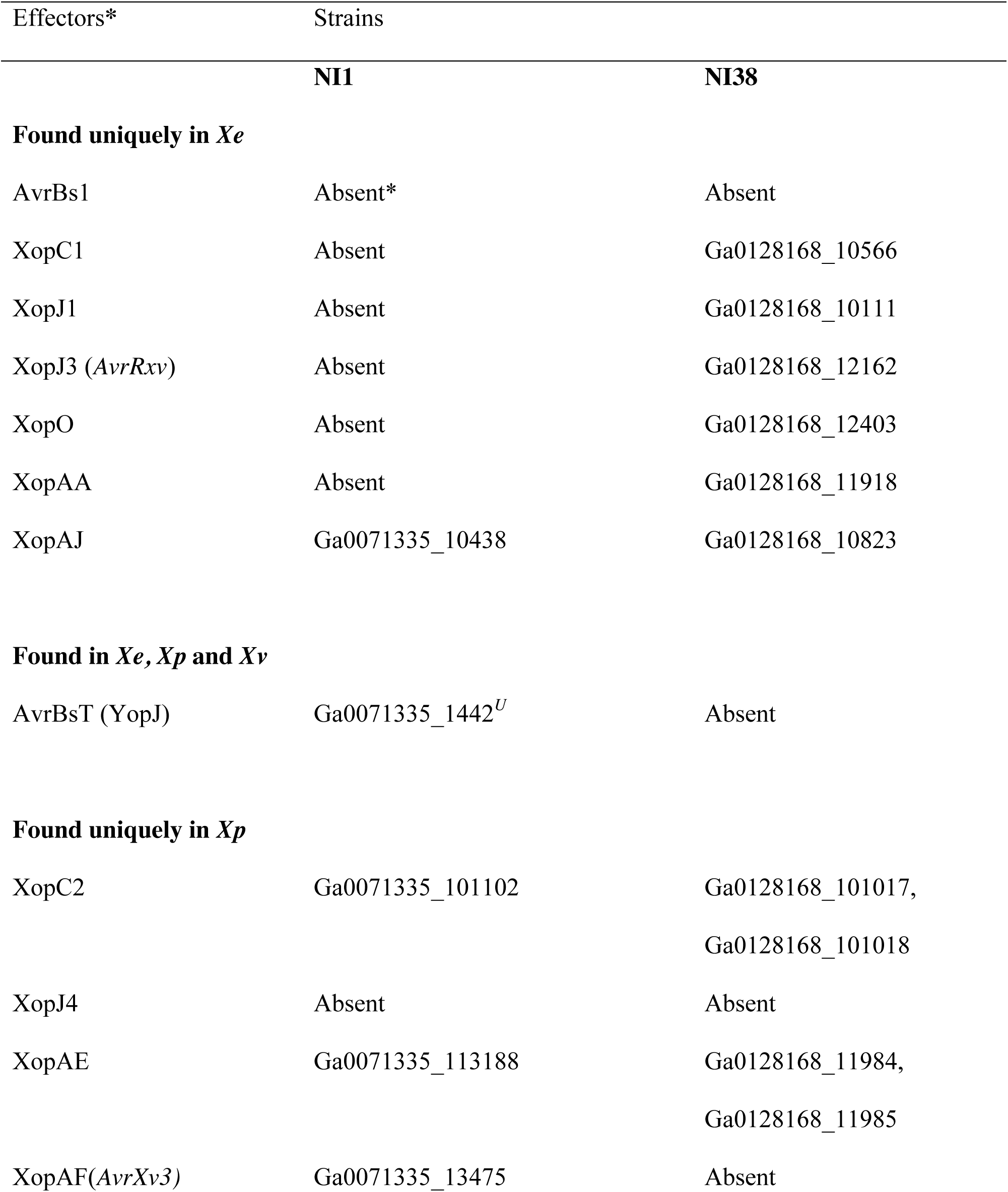
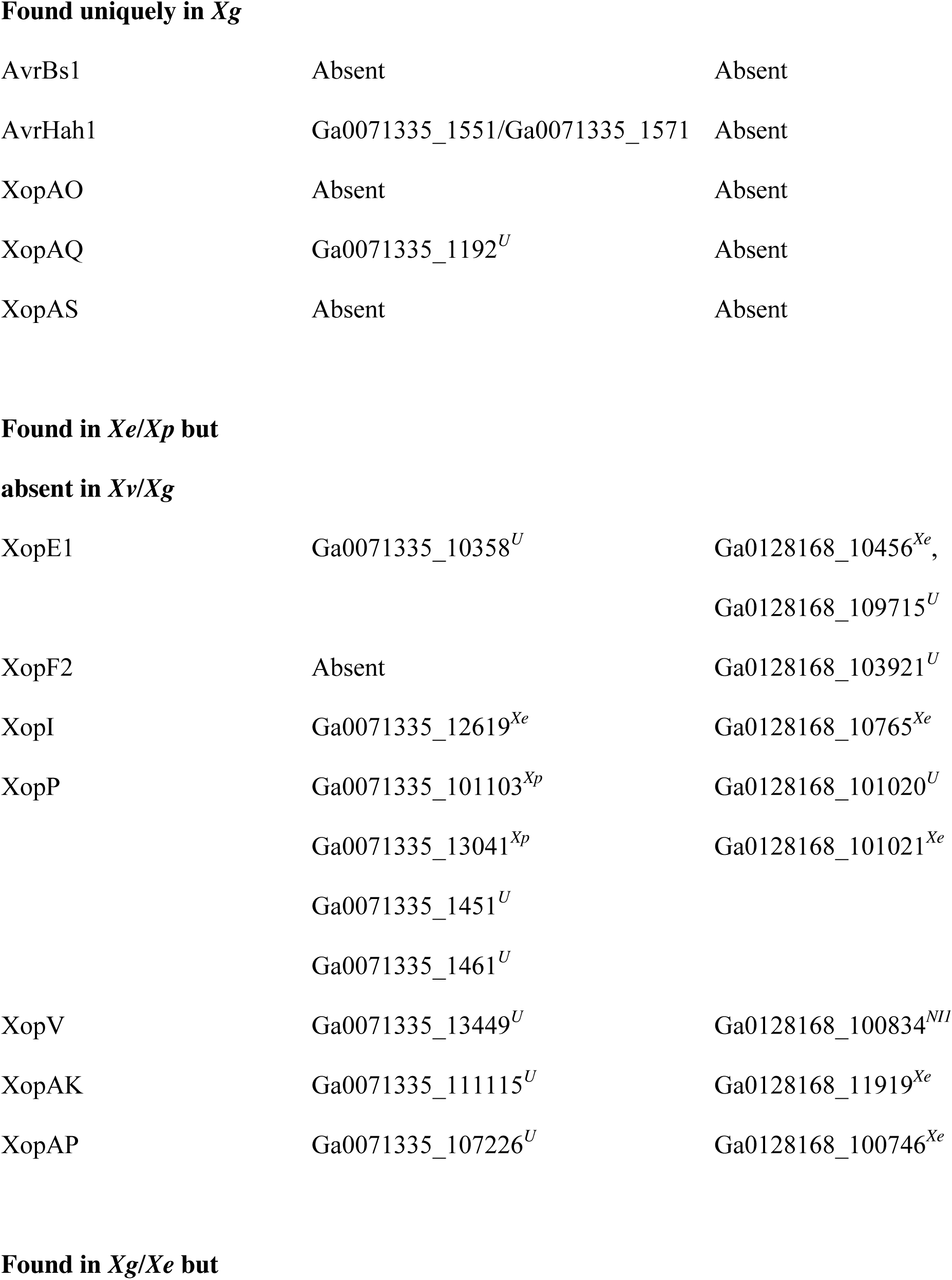
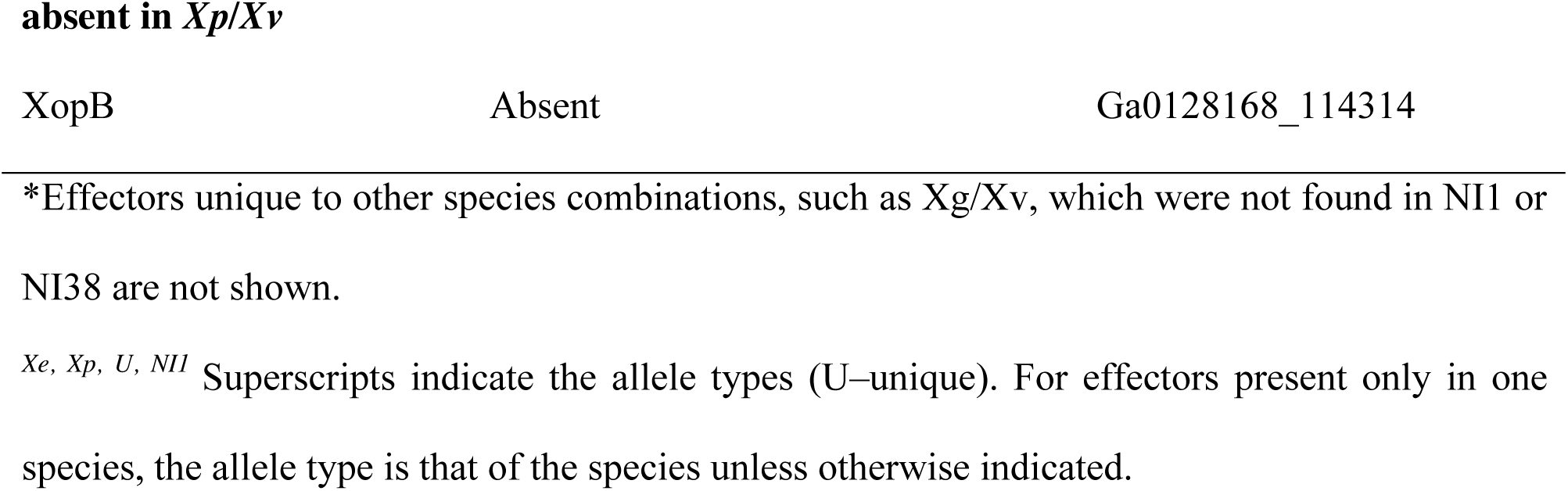
Effector profiles of NI1 and NI38 for species-specific effectors within the bacterial spot xanthomonads.

NI38 has the 11 bacterial spot core effector genes, with two copies each of *avrBs2* and *xopD*, and four copies of *xopAD* (Supplementary Table S6). NI38 has two copies each of *xopE1* and *xopP*, which are effector genes found exclusively in *Xe* and *Xp* among the bacterial spot species. Of the species-specific effectors, NI38 has all the *Xe*-specific effectors except *avrBs1*, and two copies each of *xopC2* and *xopAE* specific to *Xp*. Like other *Xe* strains, NI38 has *xopB*, which is the only effector shared by *Xe* and *Xg* with 100% amino acid identity. The *xopAJ* gene in NI38 is identical to a copy of *xopAJ* in *X. axonopodis pv. poinsenttiicola,* but a different copy of the gene from the *xopAJ* in NI1. The only *Xe* effector missing in NI38 was *xopG*, which is an effector common to *Xe*, *Xv* and *Xg*.

## Discussion

### Core genome phylogenies resolved taxonomic assignments of NI1 and NI38

Whole genome sequencing resolved the previous conflicting multilocus sequence analyses and race differentiation tests of strains NI1 and NI38, which are representative of 2014 and 2015 tomato field collections from Nigeria. We have identified these strains as novel lineages of *X. perforans* and *X. euvesicatoria,* respectively, using whole genome phylogenetic analysis. They were notably diverged from other *Xp* and *Xe* populations as a result of horizontal gene transfer and homologous recombination that has affected housekeeping genes, lipopolysaccharide clusters, secretion systems and effectors, among other genomic regions.

The close relatedness between strains of *X. euvesicatoria* and *X. perforans* is reflected in high values of average nucleotide identities in pairwise comparisons. These species were previously classified into separate species based on DNA:DNA hybridization (34) and form distinct monophyletic groups (40), but they show ANIs well above 95%. ANI identified NI1 as divergent from both *Xp* and *Xe*, and could not be used to assign this strain to either taxon.

### Recombination has shaped the evolution of X. euvesicatoria and X. perforans lineages

Our results show that recombination is a major driver of evolution of both *Xe* and *Xp*. We found evidence of both homologous and non-homologous recombination. Rates of homologous recombination varied within and between species, with *Xp* showing more evidence of recombination than *Xe*, which is perhaps most obvious in the distinct, recombinant lineages of *Xp*. Our analysis also detected higher recombination rates within the Florida population of *Xp*. The *Xe* strains were collected in the southeastern U.S., but over a longer span of time than the *Xp* strains (40). Homologous recombination is more likely to be detected in single population samples than in a timeseries (50). Recombination could be especially frequent in Florida populations, or our inclusion of single strains representing diverged lineages may have caused an underestimation of recombination in *Xp* as a whole. Additional strains from populations respresenting the diverged lineages will be required to determine if homologous recombination rates are high in other regions.

Both *Xp* and *Xe* have open and highly dynamic pangenomes, in contrast to the stability in gene composition implied when *Xanthomonas* bacterial spot pathogens are characterized by single reference genomes. Comparisons of pangenomes showed NI1 to be intermediate between *Xe* and *Xp*, whereas core genomes placed it closer to *Xp*. Our data indicate that obtaining pangenomes for *Xanthomonas* populations will be important in understanding the ongoing evolution of these pathogens.

Horizontal gene transfer and recombination likely contribute to the adaptive divergence that triggers the evolution of new bacterial spot lineages, similar to other bacterial systems such as *Bartonella henselae* and *Clostridium difficile* ST6 (51-52). The observation of extensive chromosomal replacement in NI1 was unexpected and mirrors findings in *Staphylococcus aureus* in which multiple chromosomal replacements were found in new strains compared to the reference strain, MRSA252 (52). The difference here is that NI1 was collected from a different continent than the reference strains of *X. euvesicatoria* and *X. perforans*. Nevertheless, our results for NI1 from Nigeria and 4P1S2 from Italy suggest that *X. perforans* may have very dynamic populations across the globe.

### Recombination shuffled secretion systems and effectors

Recombination has specifically affected secretion system genes and effector proteins in NI1 and NI38. Generally, characterizations of secretion systems in bacterial species have focused on reference genomes (35, 53-55), and secretions systems tend to be viewed as conserved elements of pathogenic bacteria. We found that recombination changes the gene and allelic content of secretion systems, resulting in intraspecific variation in secretion systems. The significance of the homogenization of secretion systems and effectors from two monophyletic groups in novel lineages remains to be experimentally tested. Subsequent surveys in Nigeria recovered strains like NI1 and NI38, indicating that these genotypes were stable in the short term. New combinations of effectors could be important in adaptation to hosts and/or environment, which means that recombination potentially plays an important role in the evolution of ecological interactions for these xanthomonads.

Recombination in type three secretion system genes explains why NI38 was initially identified as an *Xp* strain based on qPCR primers targeting the highly conserved *hrcN* gene. The *hrcN* allele in NI38 is distinct from the *Xp* reference strain and identical to that of 4P1S2; they both possess a single nucleotide substitution that is not present in other *Xp* strains. The nucleotide sequences where the diagnostic primers anneal in *hrcN* are identical for NI38 and all *Xp* but differ by two nucleotides from other *Xe* (73). This is the first time that the primers have not correctly differentiated *Xe* and *Xp* strains, and points to the limitations of pathogen identification using a single primer set.

The mix of type 3 secreted effectors in NI1, NI38 and 4P1S2 raises concerns regarding which effectors to use as targets for durable resistance breeding. One strategy being used to determine candidate targets for resistance breeding is to identify core, conserved effectors. Recently, comparison of sequenced genomes of *Xp* strains from Florida identified XopJ4 and AvrBST as putative stable targets for resistance breeding (39). The lack of the *xopJ4* gene in NI1 and *avrBST* in 4P1S2 makes this a less viable long-term strategy due to the possibility of introduction of these strains into Florida. Our findings suggest that bacterial spot xanthomonads may be evolving locally such that targets vary across tomato production regions, and indicate the need for global studies of effector gene content to better understand the variation in putative resistance breeding targets.

### Recombination as a cohesive and diversifying force for X. euvesicatoria and X. perforans

The close genetic distance but distinct lineages of *Xe* and *Xp* is relevant to the on-going debates on what defines a bacterial species. As observed by (7), allopatry (or at least, microallopatry) is not required for bacterial speciation, because genetic exchange rarely hinders genetic divergence in bacterial populations. Rather, recombination fosters the acquisition of novel genes and operons that aids adaptation and promotes divergence (7, 17). A competing hypothesis suggests that recombination is a cohesive force that prevents bacterial diversification and maintains lineages by homogenizing populations (56-57). Both processes may be occurring in these *Xanthomonas* populations. Thus far, most of the lineages of *Xe* and *Xp* can easily be assigned to species, while recombination is simultaneously driving the emergence of new lineages within species. However, NI1 is a more extreme case of a nearly intermediate lineage and may indicate the potential for homogenization of *Xe* and *Xp* through recombination. Both NI1 and NI38 were isolated from the same field, which is surprising because *Xp* strains are known to outcompete *Xe* strains on tomato under field conditions (58-59). NI38 strains may have acquired factors that allow co-existence with *Xp* strains on tomato.

There have been recent suggestions that both *Xe* and *Xp* be classified as a single species or as pathovars of one species (36, 60). However, these pathogens have thus far shown phenotypic differences in the lab and field, and the term pathovar implies differences in host range that is not the case with *Xe* and *Xp*. The ecotype model, which defines species by their ecology (7), may be a more appropriate concept for *Xe* and *Xp*. The ecotype model essentially defines ecotypes by their ability to coexist on the same or similar ecological resources and persist through periodic selection events. Additional studies of the genetic diversity of locally evolved lineages could provide insight into factors that drive genetic variation in effectors and other adaptive genes.

## Materials and Methods

### Sequencing of strains and calculation of average nucletide identities

After extraction of genomic DNA (61), the Nextera library preparation kit (Illumina Inc., San Diego, CA) was used to prepare genomic libraries for strains NI1 and NI38, which were subsequently sequenced using Illumina MiSeq platform. NI1 was sequenced at Kansas State University while NI38 was sequenced at the Interdisciplinary Center for Biotechnology Research, University of Florida. Draft genomes were *de novo* assembled using CLC Genomics Workbench v5, with average coverage of 259X for NI1 and 17X for NI38. The assembled sequences were annotated using the IMG/JGI platform (62). The NCBI accession numbers for NI1 and NI38 are respectively NISG00000000 and NJID00000000. Strain names and Genbank accession numbers for additional previously sequenced genomes used for this study are provided in Supplementary Table S1. The genomes of these 63 strains were previously published (63, 35-36, 40-41; 43). Pairwise Average Nucleotide Identity (ANI) based on blast was calculated using jSpecies v1.2.1 (64).

### Pan- and core genome analysis

Pan- and core genome inferences were carried out using the GET_HOMOLOGUE program (65), using default settings. All-against-all BLASTP, OrthoMCL as well as cluster of orthologous gene (COG) clustering were initially carried out to generate pan and core gene clusters before other functions within the program were used to analyze pan and core genomes (66-67). Re-sampling of genomes was performed to estimate the sizes of core and pangenomes for *X. euvesicatoria* and for *X. perforans* in the orders shown in Supplementary Table S2 (20,65). Fitted exponential decay functions were applied to resampled genomes as described by (65) and (68). A pangenome matrix (gene presence/absence) using a subset of *Xe* and *Xp* strains was generated to compare the genomes of NI1 and NI38 strains to selected *Xp* and *Xe* strains. Using the pangenome matrix, a dendrogram was constructed and a heatmap generated showing Gower’s distance using the *hcluster*_*matrix*.*sh* function in *get_homologues.pl*.

### Alignment and phylogenetic inference using core genomes

To infer the phylogeny using models for DNA sequence evolution, core genes were extracted using GET_HOMOLOGUES and filtered using an in-house python script to select for single copy orthologous genes. Alignments were carried out for each gene individually before concatenation using MUSCLE (69). We used the concatenated core genome alignment to infer a maximum likelihood phylogeny using iQTree, which was set for automatic selection of the best model (using the function –m TEST) (70). Based on the Bayesian information criterion and the corrected Akaike information criterion, the general time reversible model with proportion of invariant sites (GTR+I) was utilized as the model with the best fit out of 88 total models compared. Branch support was assessed with ultrafast bootstrap and SH-aLRT test using 1000 replicates (70). The maximum likelihood tree was visualized using FigTree v1.4.3 (http://tree.bio.ed.ac.uk/software/figtree/).

### Analysis of recombination

We carried out analysis of recombination using ClonalFrameML on *Xp* and *Xe* strains separately and together. ClonalFrameML models recombination as imports from external populations and uses a Hidden Markov Model to estimate the influence of recombination on nucleotide variation (52). For ClonalFrameML analyses, we used maximum likelihood trees generated from iQTree and core genome alignments as infiles. Transition/transversion ratios were determined by PhyML (71).

### Genome comparisons: lipopolysaccharide clusters, effectors and secretion systems

Lipopolysaccharide clusters (LPS), effectors and secretion systems were compared among strains at both the nucleotide and amino acid levels. Genes and proteins orthologous to known LPS clusters, effectors and secretion systems were identified by BLAST on the IMG platform (https://img.jgi.doe.gov/), EDGAR (edgar.computational.bio.uni-giessen.de) and from local databases (62,66,72). Secretion systems and effector allele assignments followed previous studies of *Xe* and *Xp* (35, 40).

## Acknowledgements

This work was funded in part by USDA NIFA award number 2015-51181-24312. We thank Xavier Didelot for additional help in interpreting the results of ClonalFrameML and Jose Huguet Tapia for some python scripts used in implementing get_homologues.

## Figure Legends

**Figure 1**. Pangenome comparisons based on a combined subset of *Xp* and *Xe* genomes. Values used for heatmap and dendrogram are distances between genomes based on Gower’s coefficient.

**Figure 2.** Maximum likelihood phylogeny based on concatenated core genes from of 65 genomes of *X. euvesicatoria* and *X. perforans* with midpoint rooting. Branch labels are bootstrap support values. Scale bar is in substitutions per site.

**Figure 3.** Homologous recombination in the core genomes of (A) *X. euvesicatoria* and (B) *X. perforans* estimated by ClonalFrameML. For all branches of the genealogy and positions in the genome, dark blue regions represent recombination and light blue represent invariant sites. Polymorphic sites are shown in a color indicating their level of homoplasy: white indicates no homoplasy and the range from yellow to red represent increasing degrees of homoplasy (52). The corresponding analysis of all strains together is shown in Supplementary Figure S2.

**Figure 4**. Variation in LPS clusters. Genes are colored in blocked arrows, with similar colors indicating genes belonging to the same COG (cluster of orthologous genes). a) Comparison of NI1, NI38, and reference strains for *Xtt*, *Xe* and *Xp*. b. Comparison of representative LPS clusters for *Xp* groups 1A (GEV936 and GEV1026), 1B (Xp5-6, Xp91-118), 2 (GEV1001) and the Italian strain 4P1S2. Orange arrows indicate position of *etfA* genes across the compared sets.

**Figure 5.** Reactions of a differential line carrying the *RXopJ4* gene after inoculation with a Florida *Xp* strain and strain NI1. The *Xp* strain with *avrXv4* elicited a hypersensitive response (left leaflet), while NI1 without *avrXv4* did not (right leaflet).

